# Echinocandins have an alternative mode of action on biomimetic membranes that is not directly related to the functioning of (1,3) beta-glucan synthase

**DOI:** 10.1101/2024.06.11.598481

**Authors:** Anna I. Malykhina, Svetlana S. Efimova, Ekaterina V. Vodopyanova, Natalia E. Grammatikova, Anna N. Tevyashova, Andrey E. Shchekotikhin, Olga S. Ostroumova

## Abstract

Echinocandins, one of five main classes of antifungal agents, are FDA-approved drugs for therapy of invasive candidiasis and aspergillosis. The exact mechanisms of echinocandins action are still under debates, in particular the role of target cell membranes in their function. In present study we analyzed the molecular mechanisms of action of echinocandins (anidulafungin, caspofungin, and micafungin) on the sterol-enriched lipid bilayers mimicking fungal and mammalian membranes. Calcein release assay and molecular dynamics simulations demonstrated the relation of the membrane permeability and echinocandin type. Moreover, we have shown for the first time the ability of echinocandins to form ion-permeable pores in sterol-containing membranes. Differential scanning microcalorimetry of the gel-to-liquid-crystalline lipid phase transition and confocal fluorescent microscopy of lipid domains revealed the ability of echinocandins to enhance phase segregation in membrane. Thus, echinocandins might affect the (1,3) beta-glucan synthase functioning by alteration in lipid microenvironment of target protein and cellular membrane permeability via formation of ion-selective leakage channels. We also found that small natural molecule, phloretin, potentiated the pore-forming activity of micafungin in the ergosterol-containing bilayers and significantly reduced the minimum inhibitory concentration of micafungin against fluconazole-resistant *Candida albicans, C. tropicalis*, and *C*.*krusei*.

## Introduction

Fungal pathogens are a major threat to public health as they are becoming increasingly common and resistant to treatment^1^. The invasive forms of fungal infections often affect severely ill patients and those with significant underlying immune system related conditions leading to high mortality rate^2^. In 2022 WHO published a report with a list of high priority fungal pathogens where the authors expressed concern that fungal infections receive very little attention especially in studying the antifungal resistance patterns^3^. Echinocandins, one of five main classes of antifungal drugs, are relatively new and recommended as first-line therapy by the Infectious Diseases Society of America^4^. They are highly efficient in treatment of invasive fungal infections^5^ and against fungal biofilms^6^. Echinocandins noncompetitively inhibit (1,3) beta-glucan synthase (GS) which leads to disturbance of the fungal cell wall synthesis^7^. The structure of this transmembrane protein has been recently determined by cryo-electron microscopy in two independent studies^8,9^. Both of them established that hotspots of echinocandin-resistant mutations are closely related to the membrane environment. Hu et al.^8^ noticed that hotspots were surrounded by several bound lipid molecules that alter conformation upon GS mutation, and reasoned that either echinocandin binding site lied within membrane part of GS or drugs acted through membrane changes. Zhao et al.^9^ specified that echinocandins likely inhibit glucan translocation by acting on transporting channel in membrane.Moreover, this group detected that bound lipids copurified with GS contained endogenous ergosterol which helped GS structure stabilization.

Although there is no doubt that fungicidal effect of echinocandins is related to GS inhibition, there are some non-GS related mutations that lead to echinocandin resistance. Healey et al. observed that mutations in genes disrupting sphingolipid biosynthesis resulted in caspofungin resistance in *Candida glabrata*^10^. Mutations in four different genes yielded the same changes in membrane composition: accumulation of long chain bases dihydrosphingosine and phytosphingosine. Beyond that, external addition of these substances triggered similar resistance. Furthermore, Satish et al. demonstrated that stress-induced echinocandin resistance in *Aspergillus fumigatus* brought similar membrane changes and exogenous dihydrosphingosine and phytosphingosine added to purified GS inhibited its catalytic activity^11^. Another indirect sign of membrane involvement was demonstrated in the study where caspofungin susceptibility depended on temperature in *Candida albicans* with no mutations in canonical genes^12^. There evidences together with structural findings mentioned above indicate that membrane properties are extremely important to GS proper function. The direct influence of echinocandins on target cell membranes has not been studied yet.

A recent screening study of caspofungin-sensitive strains of *Cryptococcus neoformans* has found some notable mutations that cause this sensitivity^13^. 2 out of 4 revealed deletions were in genes connected to ergosterol synthesis and led to increased sensitivity to membrane stress and elevated susceptibility to caspofungin at higher temperatures with no obvious cell wall defects. Moreover, microscopy has shown that the underlying reason behind this sensitivity was the loss of membrane integrity which contributes either by increasing the permeability to caspofungin, or by changing the location or orientation of the GS making it more susceptible to inhibition^13^. There evidences together with structural findings mentioned above indicate that membrane properties are extremely important to GS proper function. The direct influence of echinocandins on target cell membranes has not been studied yet.

Despite the fact that echinocandins are well-tolerated drugs, they have two major drawbacks that limit their use. The first one is the presence of natural resistance to these molecules in some fungus. For example, *Basidiomycetes, Mucormycetes* and *Fusarium* species exhibit resistance to echinocandins, as β-1,3-glucan constitutes only a minor component of their cell walls^14^. The second is a fast emergence of fungal isolates with acquired resistance. For instance, it took only a decade for *Candida auris*, a multidrug-resistant yeast that causes invasive infections with high mortality rate, to spread all over the world^15^. It was shown that 93% of this pathogen isolates were resistant to fluconazole, 35% to amphotericin B, and 7% to echinocandins^16^. One way to overcome the echinocandins resistance is to combine them with small molecules. Tripathi et al. showed that combination of caspofungin with a marine-derived sesquiterpene quinone, puupehenone, was highly effective against caspofungin-resistant *C. albicans, C. glabrata* and *C. neoformans*, possibly through inhibition the activity of heat shock protein 90^17^. In another study, synthetic small molecule, L-269289, was able to potentiate echinocandin activity in resistant *C. albicans, C. tropicalis* and *C. parapsilosis* by targeting geranylgeranyltransferase type I (GGTase I) and blocking the membrane localization of Rho1, the regulatory subunit of GS^18^. It seems rational to combine echinocandins with small molecules that already have antifungal activity. Thus, phloretin, a dihydrochalcone flavonoid extracted from apples, exhibits antifungal activity against *C. albicans*^19^. Proposed mechanisms of action included inhibition of the biofilm formation, suppressing the yeast-to-hyphae transition and repressing the secretion of proteases and phospholipases. Moreover, phloretin is known for its antioxidant and anti-inflammatory activity^20^. In our previous study we showed that phloretin demonstrates dramatic membrane activity that implemented via changing in membrane electrical and elastic properties and led to significant increase in pore formation of polyene antifungals, amphotericin B and nystatin^21,22^.

Formation of biofilms by many pathogenic fungi significantly complicates antifungal treatment especially for immunocompromised patients^23^. For example, *Candida spp*. are now the third most common microorganisms associated with catheter-related blood stream infections^24^. Encapsulation of antifungals into liposome formulation might improve their antifungal activity, drug solubility, reduce toxicity, and overcome resistance. For instance, more efficacious penetration of the fluconazole liposomal form resulted in growth inhibition of resistant biofilm-forming *C. albicans* isolates^25^. Liposomal amphotericin B was mainly developed to resolve high toxicity which is known for this drug, additionally, a rapid accumulation at the infection site (including biofilms) was observed for this formulation^26^. To the date, only anidulafungin among echinocandins was studied in liposome nanoparticles^27^. Although a minimum inhibitory concentration against *C. albicans* was equivalent to free drug, liposomal formulation exhibited superior biofilm disruption reducing fungal burden by 99%. Thus, liposomal form of echinocandins could overcome the fungal resistance associated with biofilms.

In our work we implemented complex approach to investigate the action of echinocandins, anidulafungin (ANF), caspofungin (CSF), and micafungin (MCF) (Figure 1), on the model lipid membranes, including fluorescence assay for calcein leakage from lipid vesicles, registration of ionic currents through planar lipid bilayers, differential scanning microcalorimetry of membrane lipid phase transition, confocal fluorescence microscopy of lipid domains in liposomes, and molecular dynamics simulations. The main findings concerned the induction of calcein release, the formation of transmembrane pores of weak cation/anion selectivity, and the enhancement of lipid phase segregation. The effects depended on echinocandin type and sterol composition of target membranes. Moreover, plant polyphenol phloretin was found to potentiate the pore-forming ability of micafungin in ergosterol-enriched bilayers. The potential effect of combining micafungin with phloretin was assessed against *Candida* spp. Thus, our results revealed a possible alternative antifungal mechanism of action of echinocandins that might be related to alteration in lipid microenvironment of GS and increase in membrane permeability. A combination of micafungin with phloretin seems to be a promising approach to increase the antibiotic efficacy by alternative mode of action.

**Figure 1.**
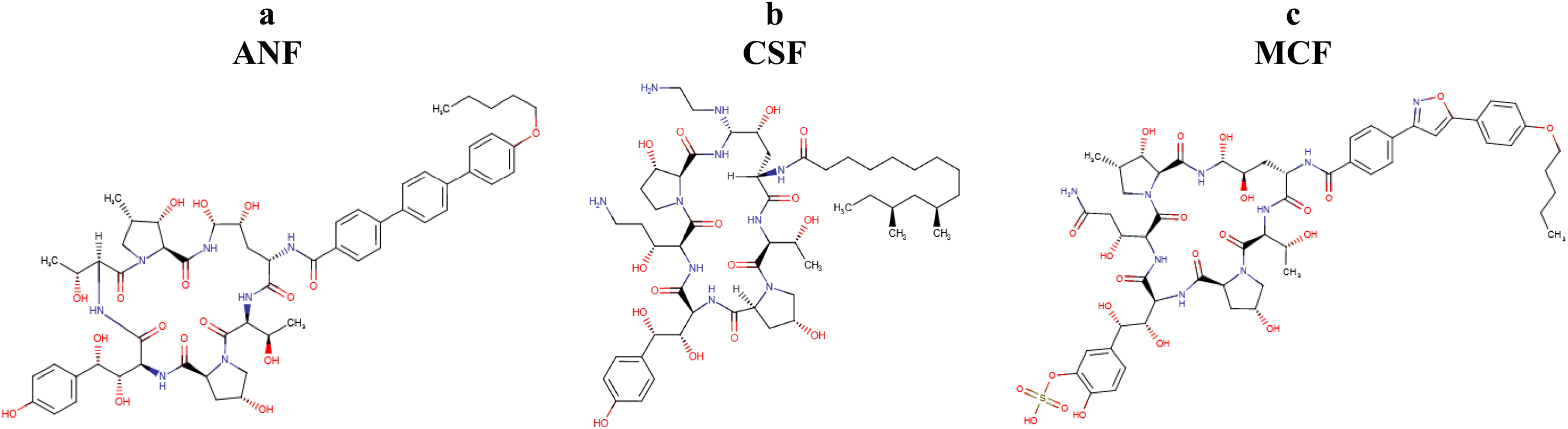
Chemical structures of tested echinocandins: (**a**) anidulafungin (ANF), (**b**) caspofungin (CSF), (**c**) micafungin (MCF).

## Materials and methods

Experimental procedures, materials and methods are provided in Supplementary material.

## Results and discussion

### Assessment of permeability of model lipid membranes produced by echinocandins

The interaction of echinocandins, ANF, CSF, and MCF, with membranes was tested on lipid unilamellar vesicles prepared from POPC admixed with ergosterol (67:33 mol.%) or cholesterol (67:33 mol.%). The lipid content was chosen in order to imitate fungal and mammalian membranes accordingly^28-31^. To assess contribution of sterols in this interplay, vesicles composed of pure POPC content were tested separately. Figure 2 presents the results obtained: ANF led to calcein efflux from liposomes of all tested compositions, CSF was active in pure POPC and POPC:Erg (67:33 mol.%) vesicles, while MCF didn’t show any significant activity in all lipid models. The kinetic characteristics of the calcein release from liposomes are presented in Table 1. Characteristic time of calcein leakage, τ, from POPC, POPC:Chol (67:33 mol.%), POPC:Erg (67:33 mol.%) liposomes induced by ANF was equal to 14, 6, and 15 min, respectively. While addition of CSF into suspension of POPC and POPC:Erg (67:33 mol.%) vesicles produced fluorescent marker release with characteristic times of 147 and 125 min, respectively (Table 1). *IF*_*max*_ in the presence of ANF decreased in the row of POPC (about 85%) ≥ POPC:Erg (about 73%) > POPC:Chol (47%). The lipid selectivity of CSF expressed in *IF*_*max*_ values diminished in the following order: POPC ≈ POPC:Erg (about 75%) >> POPC:Chol (1%). Thus, both echinocandins, ANF and CSF, were more active in pure POPC and Erg-enriched membranes compared to Chol-containing bilayers. As it was noted above, MCF did not disengage calcein from liposomes of all tested compositions: *IF*_*max*_ was not more than 1%. Analyzing the presented data, we hypothesized that ANF was able to form transmembrane defects in liposomes of various composition and enforced quick calcein leakage through these channels. MCF was not characterized by the ability to form calcein-permeable transmembrane defects in all tested lipid models. The slow CSF-induced disengagement of fluorescent marker from pure POPC and Erg-containing vesicles might be addressed to disintegration of liposomal membranes rather than the formation of calcein-permeable pores.

**Table 1.** The parameters characterizing the dependences of echinocandin-induced calcein leakage on time

| parameter | POPC |  |  | POPC:Chol<br>(67:33 mol.%) |  |  | POPC:Erg<br>(67:33 mol.%) |  |  |
| --- | --- | --- | --- | --- | --- | --- | --- | --- | --- |
|  | *ANF* | *CSF* | *MCF* | *ANF* | *CSF* | *MCF* | *ANF* | *CSF* | *MCF* |
| $IF_{max}$, % | 85 ± 5 | 74 ± 5 | 1 ± 1 | 47 ± 4 | 1 ± 1 | 1 ± 1 | 73 ± 4 | 76 ± 1 | 1 ± 1 |
| $\tau$, min | 14 ± 5 | 147 ± 34 | N/A | 6 ± 2 | N/A | N/A | 15 ± 5 | 125 ± 61 | N/A |
| $R^2$ | 0.9 | 0.9 | N/A | 0.8 | N/A | N/A | 0.9 | 0.9 | N/A |

$IF_{max}$ – maximal leakage of calcein from liposomes, $\tau$ – the characteristic time of one-exponential dependence fitting the time dependence of calcein release from liposome, $R^2$ – correlation coefficient, N/A – not applicable.

**Figure 2.**
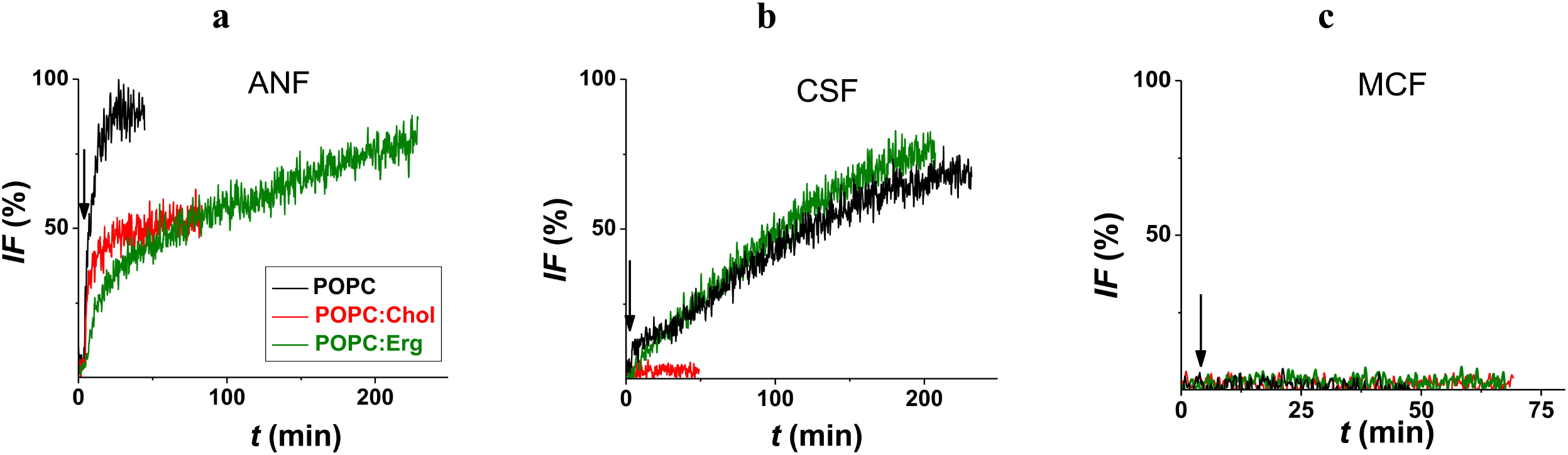
Time-dependence of relative fluorescence of calcein (*IF*, %) leaked from vesicles composed of POPC (*black curves*), POPC:Chol (67:33 mol.%) (*red curves*), and POPC:Erg (67:33 mol.%) (*green curves*). The moments of addition of ANF (**a**), CSF (**b**), MCF (**c**) into a liposomal suspension up to 50 μM are indicated by arrows.

We also examined the ability of echinocandins to increase ion permeability of lipid bilayers. The addition of ANF, CSF, and MCF at *cis*-side of the bilayer up to 1-2 μM in the membrane-bathing solution led to the appearance of step-like current fluctuations of various amplitudes in the picoampère range through Erg-containing bilayers in an asymmetric salt system (under conditions with different electrolyte concentrations in aqueous solutions on different sides of the membranes). Inset in Figure 3 shows typical current traces demonstrating ANF (a), CSF (b), and MCF (c) pores in the POPC:Erg (67:33 mol.%) membranes at different transmembrane voltages. Replacement of Erg for Chol in the membrane composition did not affect pore-forming activity of echinocandins (Figure S1, insets, Supporting Information).

**Figure 3.**
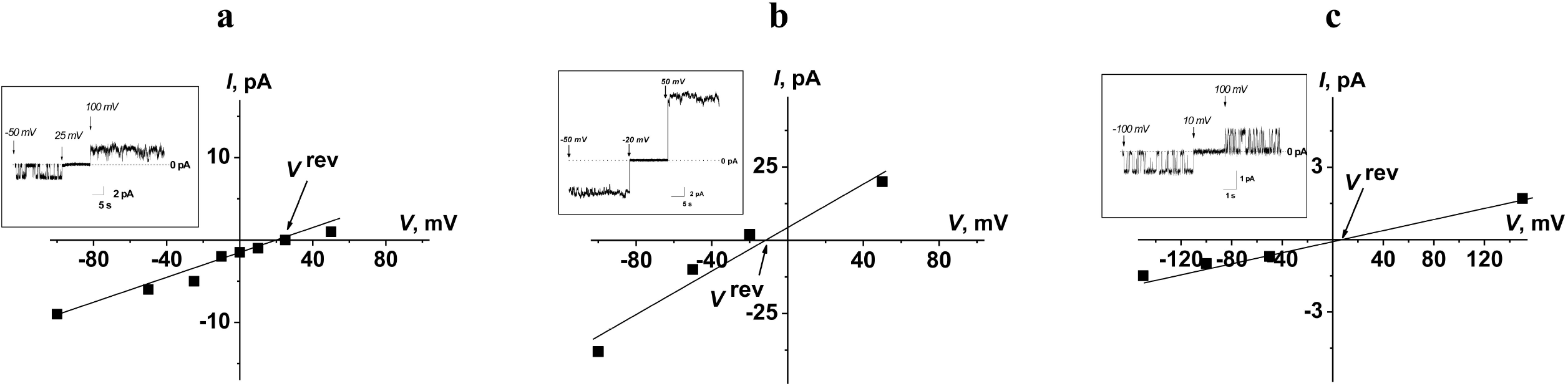
Cation/anion selectivity of transmembrane pores induced by ANF (**a**), CSF (**b**) and MCF (**c**). *I–V* curves of modified with sufficient amount of echinocandins (cis-side only) to induce a bilayer permeability. The membranes were composed of POPC:Erg (67:33 mol.%) and bathed in asymmetric salt solutions of 0.025 M NaCl (10 mM HEPES, pH 7.4 (*cis*-side)) and 0.15 M NaCl (10 mM HEPES, pH 7.4 (*trans*-side)). Arrows indicate the reversal potentials (*V*_*rev*_). *Inset*: Current fluctuations of the ANF (**a**), CSF (**b**) and MCF (**c**) pores at different transmembrane voltages in asymmetric solutions.

The cation/anion selectivity of ANF and CSF channels in POPC:Chol (67:33 mol.%) and POPC:Erg (67:33 mol.%) bilayers was determined. A potential of zero current, *V*^*rev*^, was measured after the bilayer modified with sufficient only *cis-*side ANF or CSF addition (Figure 3). Figure 3 demonstrates the different cation/anion selectivity of the bilayers treated with ANF, CSF and MCF in the Erg-containing bilayers. The values of the potential corresponding to zero current, *V*^*rev*^, in the presence of ANF, CSF and MCF were equal to 25 ± 5, –12 ± 7, and 9 ± 5 mV, respectively. This reversal potential corresponded to the cation (Na^+^) transfer numbers, equal to 0.7 ± 0.1, 0.4 ± 0.1, and 0.6 ± 0.1 for ANF, CSF, and MCF, respectively. The cation transfer numbers in the Chol-containing membranes were equal to 0.8 ± 0.1, 0.4 ± 0.1, and 0.6 ± 0.1 for ANF, CSF, and MCF, respectively (Figure S1, Supporting Information). This result clearly indicated that the ability of echinocandins to form ion-permeable pores in the membranes did not significantly depend on their sterol composition. The most probable explanation for preferential cation selectivity of ANF and MCF pores is related to partial negative charges on carbonyl groups in ANF and MCF rings. An extremely weak anion selectivity of CSF pores might be explained by the presence of protonated amino groups in side radicals of CSF. Also, it can be proposed that the sterol molecules are weakly involved in the pore-formation by echinocandins. The lack of correlation between the abilities of echinocandins to cause fluorescent marker leakage from liposomes (Table 1) and to form ion-selective pores in planar lipid bilayers (Figure 3, Figure S1, Supporting Information) did not allow us to directly link these effects.

Molecular dynamic simulation and umbrella sampling method were employed for assessment of membrane permeability for echinocandins. The potential of mean force represents the relative free energies of a given molecule at different distances from membrane center along Z-axis (Figure 4). During the initial movement of the echinocandin from the solution to headgroup region of the lipid bilayer, there was a slight decrease in free energy (no more than 1 kcal/mol) which reflected favorable electrostatic interaction between polar group of the molecule and the membrane (Figure 4, Table 2). Subsequently, the free energy gradually increased as the echinocandin molecule passed through membrane with the maximum value at the center of hydrophobic core. The dissimilarity between membranes as well as between compounds was observed. For all echinocandins the lowest energy difference was for pure POPC membrane (ΔG for ANF, CSF and MCF was equal to 24.9 kcal/mol, 28.3 kcal/mol, and 36.1 kcal/mol, respectively) (Table 2). Sterol inclusion into membrane composition made the membrane thicker and almost twice less permeable for tested molecules. The energy differences for Chol-containing membrane were 42.5 kcal/mol, 50.0 kcal/mol, and 52.3 kcal/mol, and for Erg-enriched bilayer – 39.4 kcal/mol, 41.2 kcal/mol, and 55.7 kcal/mol for ANF, CSF and MCF, respectively (Table 2). The distinction between sterols was minimal (≈ 3 kcal/mol) for ANF and MCF, while for CSF it was surprisingly large (≈ 9 kcal/mol). The presented data clearly demonstrate that the penetration of echinocandins through the membranes is characterized by a large energy barrier due to high costs of molecule transport from the polar microenvironment to the center of the lipid bilayer. It is worth noting that the data of umbrella sampling (Figure 4, Table 2) were consistent with the results of release measurements (Figure 2, Table 1). In particular, MCF that did not cause calcein leakage (Figure 2, Table 1) was characterized by the highest energy of entry into the center of the bilayer (Figure 4, Table 2). The energies of ANF and CSF translocation through membranes made of pure POPC or its mixture with Erg were lower than through Chol-enriched bilayers (Figure 4, Table 2), which coincides with the greater leakage of the marker from POPC liposomes and Erg-containing vesicles compared to Chol-enriched liposomes (Figure 2, Table 1). Moreover, there was the more pronounced sterol dependence of the free energy of CSF translocation than of ANF (Figure 4, Table 2) similar to difference in IF_max_ values in the presence of these echinocandins (Figure 2, Table 1). Thus, echinocandin-induced calcein leakage might be associated not with the formation of pores in membranes, but with the ability of antibiotics to penetrate the bilayer.

**Table 2.** Parameters charactering the echinocandin action on the membranes of various composition in the molecular simulation

| agent | POPC |  | POPC:Chol<br>(67:33 mol.%) |  | POPC:Erg<br>(67:33 mol.%) |  | *N* | *logP* |
| --- | --- | --- | --- | --- | --- | --- | --- | --- |
| | $\Delta G$,<br>kcal/mol | *APL*, Å<sup>2</sup> | $\Delta G$,<br>kcal/mol | *APL*, Å<sup>2</sup> | $\Delta G$,<br>kcal/mol | *APL*, Å<sup>2</sup> | | |
| *control* | – | 63.4 ± 1.4 | – | 45.1 ± 1.2 | – | 46.7 ± 1.1 | – | – |
| *ANF* | 24.9 | 64.0 ± 1.4 | 42.5 | 45.9 ± 1.3 | 39.4 | 47.2 ± 1.1 | 7.5 | -1.5 |
| *CSF* | 28.3 | 64.8 ± 1.5 | 50.0 | 46.7 ± 1.2 | 41.2 | 46.7 ± 1.1 | 8.0 | -4.8 |
| *MCF* | 36.1 | 64.0 ± 1.3 | 52.3 | 46.2 ± 1.4 | 55.7 | 46.9 ± 1.4 | 9.8 | -6.3 |

$\Delta G$ – the difference in potential of mean force between membrane center (0 nm) and water (3nm); *APL* – area per lipids; *N* – number of hydrogen bonds formed between echinocandin molecule and water; *LogP* values (logarithms of the octanol/water partition coefficients) were predicted using [https://go.drugbank.com/].

**Figure 4.**
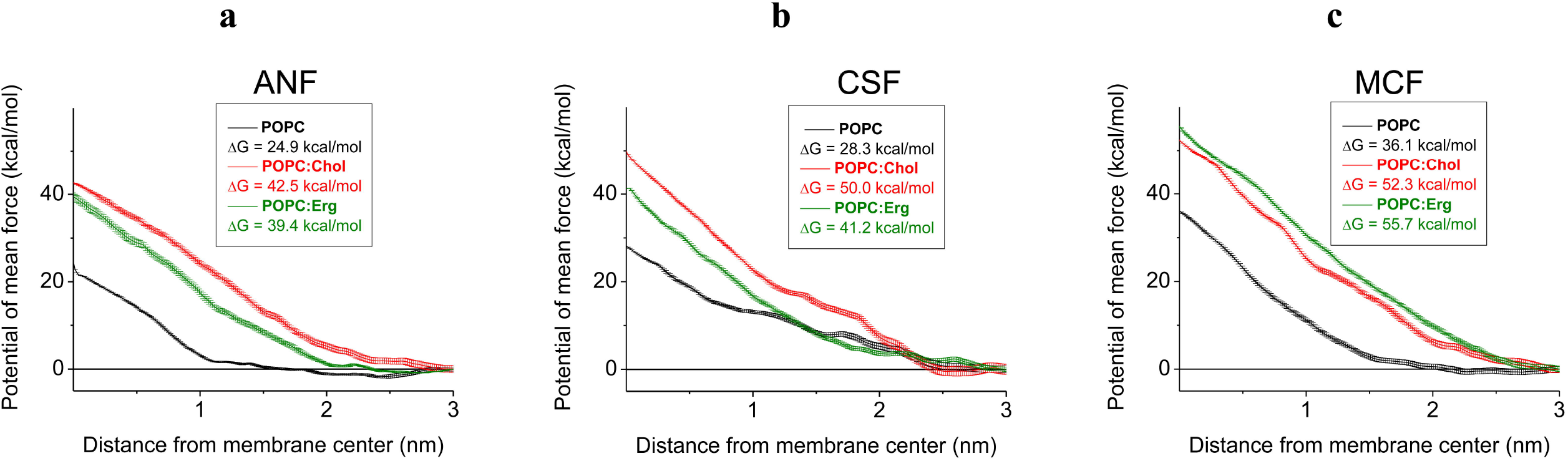
The potential of mean force is plotted for three model membranes containing pure POPC (*black curves*), POPC:Chol (67:33 mol.%) (*red curves*), POPC:Erg (67:33 mol.%) (*green curves*) for each echinocandin molecule separately: ANF(**a**), CSF (**b**), and MCF (**c**). The free energy value was set to zero in solvent region (3 nm from membrane center). Error bars represent the standard error calculated by the Bootstrap method.

The observed dependence might be related to hydrophilic properties of the echinocandin molecules. According to the literature data *logP* is equal to -1.5, -4.8, -6.3 for ANF, CSF and MCF, respectively (Table 2). To demonstrate the relation, we performed molecular dynamic simulation of one echinocandin molecule in solution for 100 ns and then assessed the number of hydrogen bonds formed between the substance and the water molecules. Each echinocandin molecule has groups that could act as donors or acceptors. We observed that average number of all hydrogen bonds also increased in a row of ANF < CSF < MCF and composed 7.5, 8.0 and 9.8 bonds accordingly (Figure S2, Supporting Information, Table 2). In interaction with water echinocandins act mostly as acceptors. Thus, the higher polarity of MCF among tested echinocandins is in a good agreement to its lower membrane permeability.

In order to try to unravel the reason for more pronounced sterol dependence of CSF-induced calcein release and Δ*G* of its translocation compared to other tested echinocandins (Table 2), the detailed analysis of hydrogen bonds of CSF in sterol-containing membranes during steered molecular dynamic was performed (Figure S3, Supporting Information). The analysis revealed that in the POPC:Chol (67:33 mol.%) membrane CSF had tendency to remain being bound by hydrogen bonds longer than in Erg-enriched bilayer. Further review showed that the main contribution to this lagging was introduced by water. Analyzing Table 2, one can notice that CSF was able to produce an increase in APL in Chol-containing membranes, while it did not affect APL in Erg-enriched bilayers. The increase in area might help to explain the ability of CSF to pass along with water, whereas the formation of this hydrophilic pathway may require additional energy.

### Modifying of lipid phase organization by echinocandins

Differential scanning microcalorimetry of giant unilamellar vesicles was used to investigate the lipid phase transitions. In the absence of echinocandins the melting temperature of DPPC, DPPC:Chol (85:15 mol.%) and DPPC:Erg (85:15 mol.%), *T*_*m*_, was equal to 41.5 ± 0.1ºC, 40.6 ± 0.2ºC, and 40.5 ± 0.1 ºC, respectively. The width of the peak related to main transition of DPPC, DPPC:Chol (85:15 mol.%) and DPPC:Erg (85:15 mol.%), Δ*T*_*b*_, was equal to 2.6 ± 0.3ºC, 3.8 ± 0.2ºC, and 4.9 ± 0.2°C, respectively. Tested echinocandins were shown to strongly affect the thermotropic behavior of pure DPPC and its mixtures with different sterols. Figure 5 shows the representative heating thermograms of DPPC, DPPC:Chol (85:15 mol.%) and DPPC:Erg (85:15 mol.%) vesicles before and after addition of echinocandins into liposome suspension up to various lipid:echinocandin molar ratios. ANF, CSF and MCF completely abolished the gel-to-ripple phase transition of pure DPPC (Figure 5) which was observed at 35.1 ± 0.5ºC in the absence of antibiotics. The molecular origin of ripple-phase formation is associated with the DPPC headgroup region and adsorption of echinocandins might upset the balance between lipid heads and chains. The main peaks on thermograms in the presence of echinocandins (at least at low lipid:antibiotic molar ratio) acquired a complex profile, characterized by the presence of two or three overlapping components (Figure 5). This indicated the existence of several lipid phases enriched with different concentration of echinocandins. Table S1 (Supporting Information) presents the results of decomposition/deconvolution analysis of main DPPC, DPPC:Chol (85:15 mol.%) and DPPC:Erg (85:15 mol.%) transition peaks in the presence of echinocandins. To assess the dependences of lipid thermotropic behavior on echinocandin type, mean melting temperature, *T*_*m_mean*_, was determined according to eq. 3 (Material and Methods section). Addition of ANF and MCF caused a dose-dependent decrease in mean melting temperature, *T*_*m_mean*_, and increase in the width of the transition, Δ*T*_*b*_, of DPPC, DPPC:Chol, and DPPC:Erg, while CSF did not practically affect these parameters (Figure 6). Moreover, ANF demonstrated notable lipid specificity of action: it had a more pronounced effect on melting temperature of sterol-enriched membranes than pure DPPC (Figure 8, upper panel). The ability of MCF to decrease *T*_*m_mean*_ did not distinctly depend on the type of lipid model (Figure 6, upper panel). A decrease in the lipid melting point and a decrease in the sharpness of the phase transition in the presence of ANF and MCF indicated their disordering effect on membrane-forming lipids. ANF and MCF might affect lipid melting through immersion into the membrane and increase in APL.

**Figure 5.**
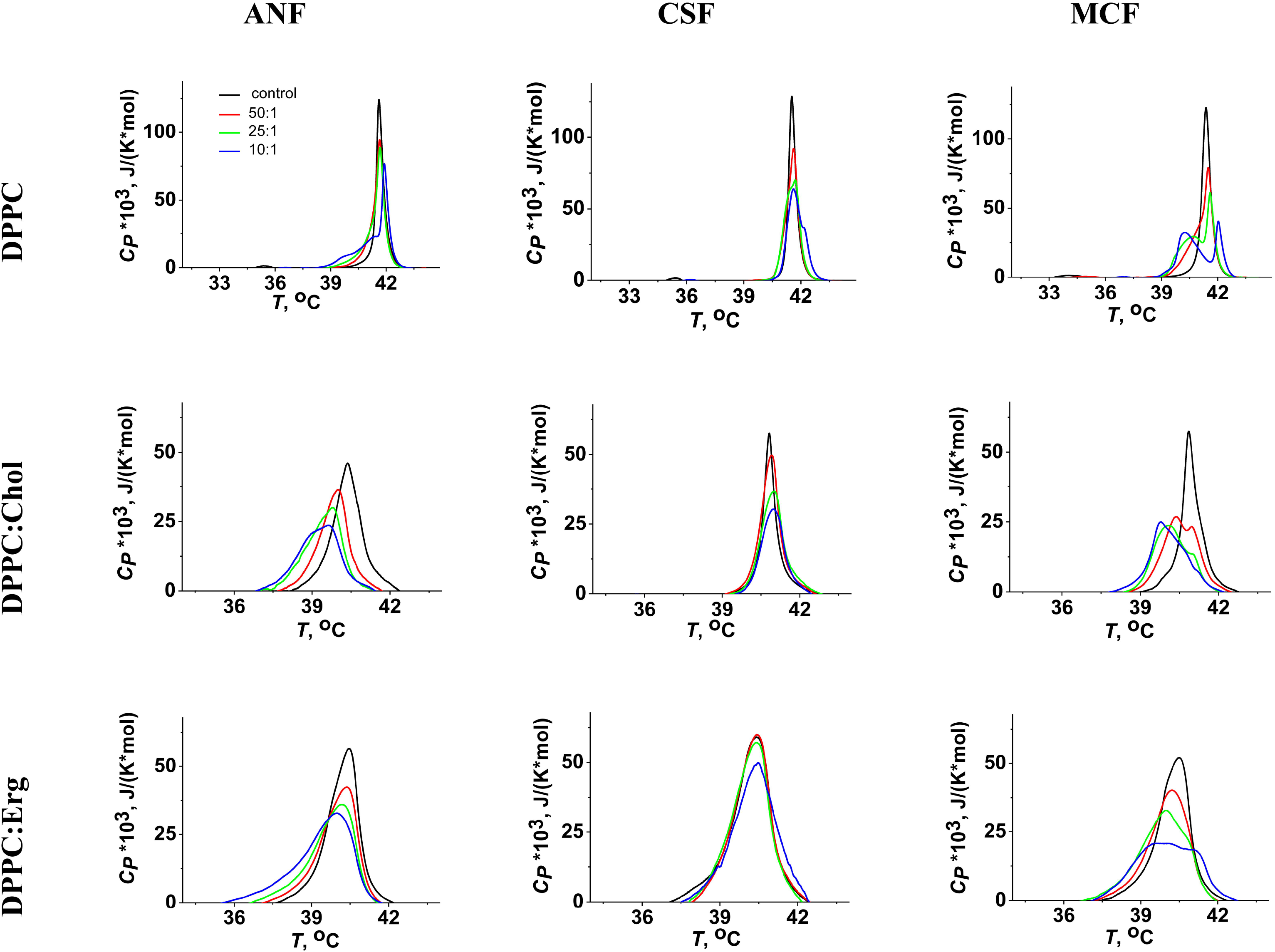
Heating thermograms of DPPC (*top panel*), DPPC:Chol (85:15 mol.%) (*medium panel*) and DPPC:Erg (85:15 mol.%) (*bottom panel*) liposomes in the absence (control, *black curves*) and presence of ANF (*left column*), CSF (*medium column*) and MCF (*right column*) at lipid:echinocandin ratio of 50:1 (*red curves*), 25:1 (*green curves*), and 10:1 (*blue curves*).

**Figure 6.**
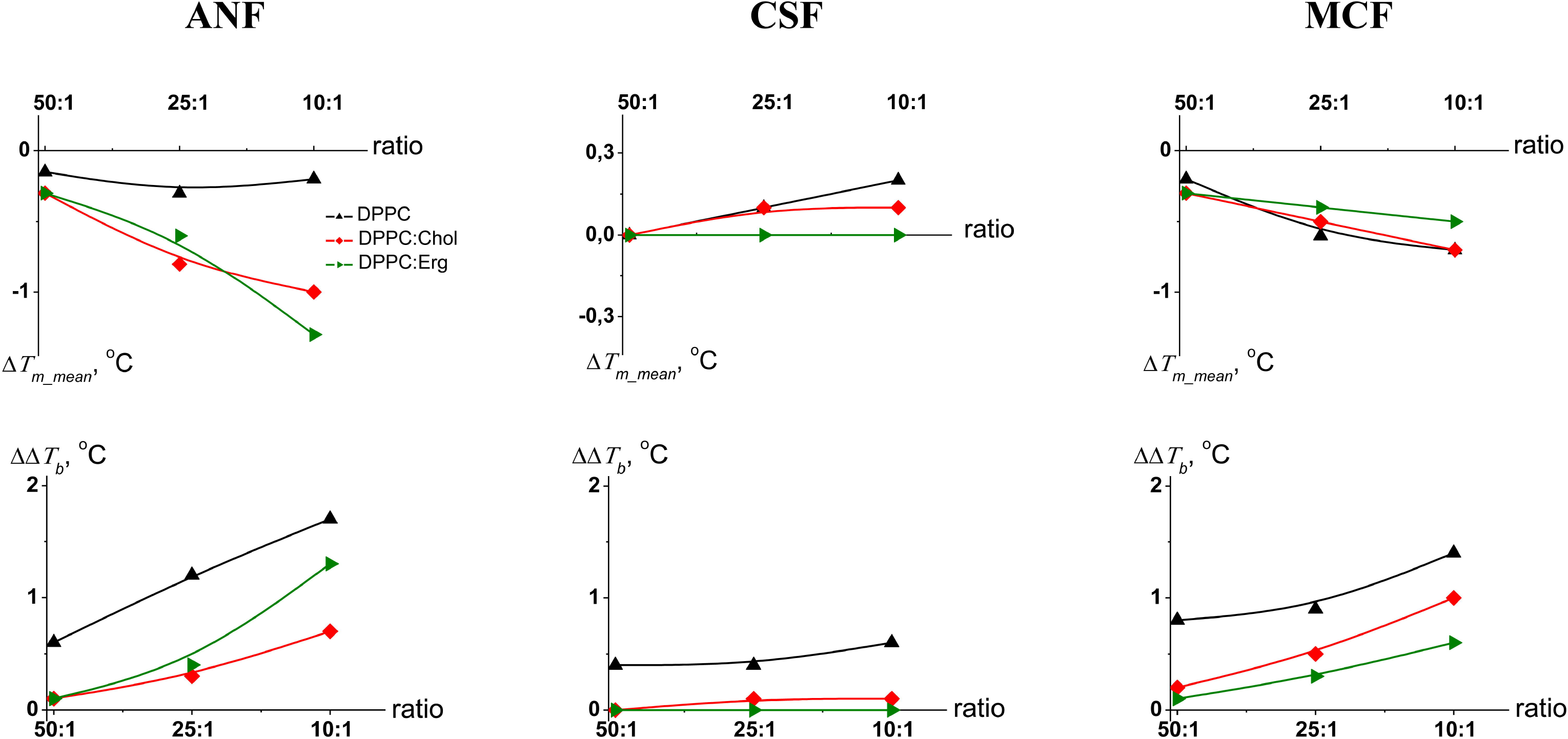
Dependence of the parameters characterizing the thermotropic behavior of DPPC (*black curves*), DPPC:Chol (85:15 mol.%) (*red curves*) and DPPC:Erg (85:15 mol.%) (*green curves*) on lipid:echinocandin molar ratio in the presence of ANF (*left column*), CSF (*medium column*) and MCF (*right column*). The changes in mean melting temperature of heating scan (Δ*T*_*m_mean*_) and a width of the main peak (ΔΔ*T*_*b*_) are presented on top and bottom panels respectively.

It was not possible to directly compare the DSC results with the molecular simulation data presented in Table 2, where APL was assessed in 100 ns molecular simulation with one molecule of the drug inserted in the model membrane. ANF- and MCF-induced increase in APL value in POPC, POPC:Chol, and POPC:Erg bilayers (Table 2) were in accordance with differential scanning microcalorimetry results demonstrating disordering action of ANF and MCF (Figure 5, Figure 6). Wherein, CSF having insignificant effects on lipid melting (Figure 5, Figure 6) showed more pronounced increase in APL in POPC and POPC:Chol membranes, and no action on APL in the POPC:Erg bilayer (Table 2). This discrepancy was not related to phospholipid type, because CSF increased APL in both POPC (Table 2) and DPPC membrane (data not shown). A possible explanation might be related to cooperativity effect of echinocandins and especially CSF on lipid phase behavior. Supporting evidence for such explanation may be taken from another molecular simulation where one echinocandin molecule was placed in solution near membrane. ANF and MCF but not CSF spontaneously imbedded in membrane (data not shown). Indeed, CSF due to the absence of polar groups in the tail, interacted with membrane only by head, thus, we could anticipate cooperativity of CSF molecules in real experiment for membrane incorporation.

Based on differential scanning microcalorimetry data obtained, we proposed that tested echinocandins might influence the phase separation in the membrane. Therefore, lipid domain organization in GUV membranes in the presence of ANF, CSF and MCF was assessed by confocal fluorescence microscopy (Figure 7). Vast majority of DOPC:Chol (80:20 mol.%) GUVs possessed liquid disordering phase in both control experiments with untreated liposomes and after echinocandin addition (Figure 7a). Figure 7b shows typical micrographs of homogeneously colored GUVs. Few liposomes with multiple round domains that should be associated to liquid ordered phase were detected in all samples with no difference between conditions (Figure 7c). However, echinocandins had a peculiar effect on domain morphology: they formed homogeneously colored GUVs (Figure 7d) and quasi gel phase visually presented as dendritic domains of uneven shape (Figure 7e). We speculated that adsorption of echinocandin molecules having large polar heads and only one hyrophobic tail at the interface between ordered and disordered lipid phases should diminish hydrophobic mismatch by reducing surface tension. It is known that GS is embedded in lipid raft^11^ and the reorganization of the lipid domains by echinocandins could also violate GS function.

**Figure 7.**
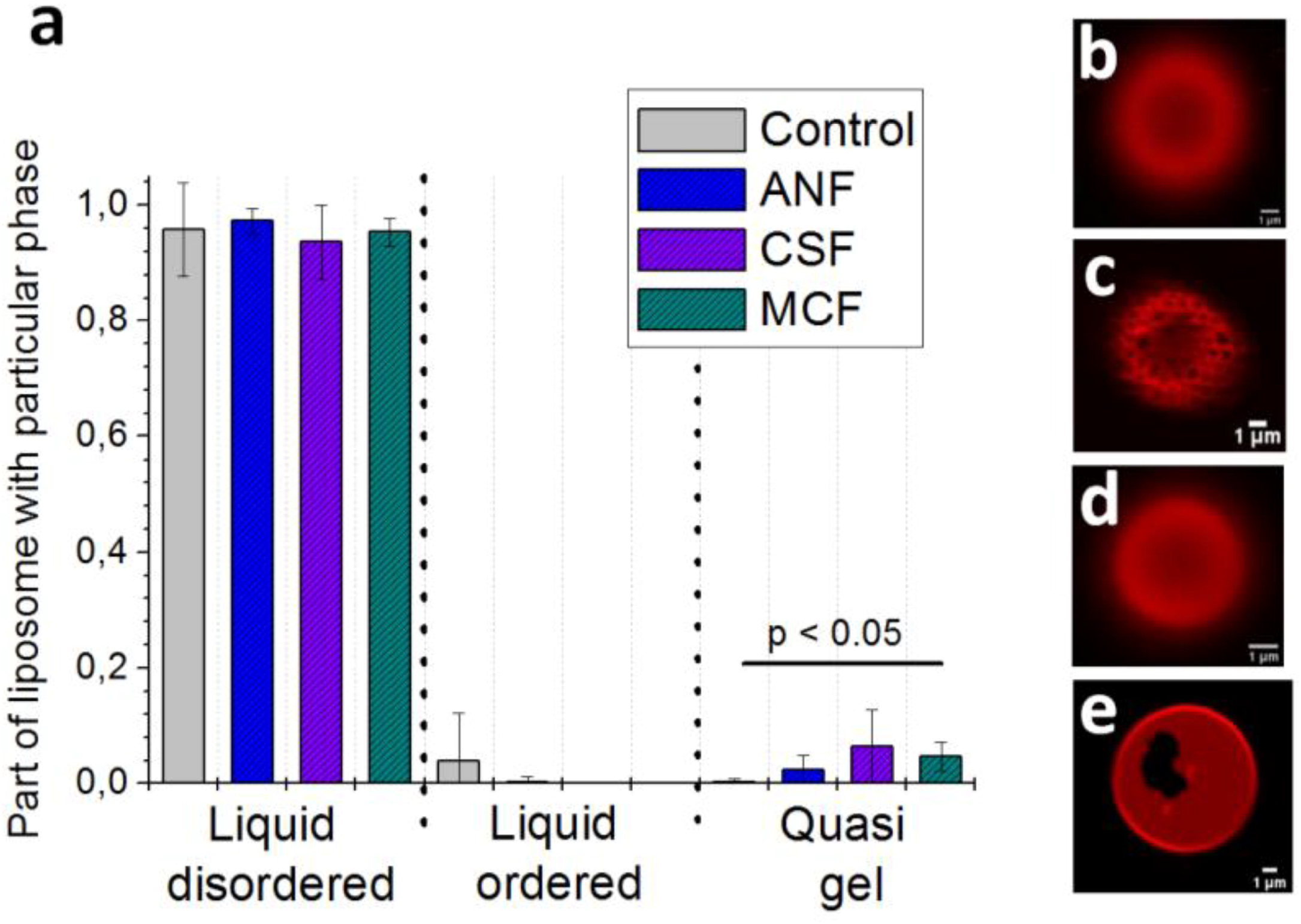
(**a**) Bar chart of liposome distribution with particular lipid domains (liquid disordered, liquid ordered and quasi gel) in control (light green color) and after addition of 50 *µ*M ANF (*blue color*), 10 *µ*M CSF (*violet color*), and 50 *µ*M MCF (*dark green color*). The difference in uncolored lipid domain morphology between conditions was statistically significant (*p*-value < 0.05, ANOVA). (**b-e**) Typical GUVs confocal micrographs in control (**b, c**) and in the presence of MCF (**d**,**e**)(scale bar is equal to 1 µm) possessing liquid disordered phase (**b, d**), liquid ordered (**c**), quasi gel (**e**) domains.

**Figure 8.**
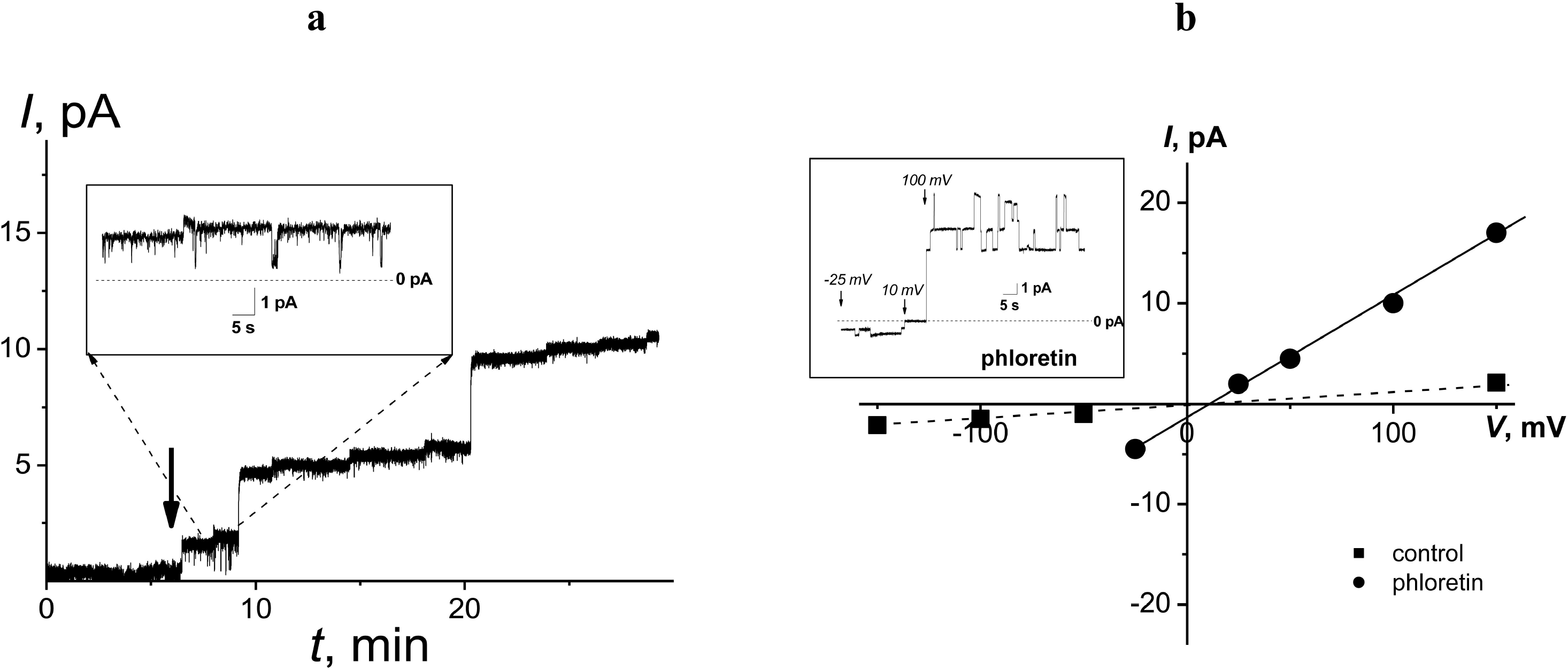
(**a**) The effect of addition of phloretin in the bathing solution at both-side of membrane up 20 μM on the steady-state bilayer conductance induced by *cis*-side addition of 5 µM MCF. MCF was added at the initial moment. Arrow indicates the moment of introduction of phloretin to the bilayer bathing solution. *V* = 100 mV. *Inset*: Current fluctuations of the MCF pores in asymmetric solutions after addition of phloretin. (**b**) *I–V* curves of bilayer modified with MCF (*cis*-side only) before (dotted line) and after phloretin addition (solid line). The membrane was composed of POPC:Erg (67:33 mol.%) and bathed in asymmetric salt solutions of 0.025 M NaCl (10 mM HEPES, pH 7.4 (cis-side)) and 0.15 M NaCl (10 mM HEPES, pH 7.4 (trans-side)). *Inset*: Current fluctuations of the MCF pores at different transmembrane voltages in asymmetric solutions after addition of phloretin.

### Potentiation of pore-forming ability of micafungin by phloretin

According to the literature data MCF is a drug that was especially designed to have minimal disruption of red blood cells^32^. For this reason, we tried to enhance pore-forming activity of this echinocandin by using plant-derived small molecule, phloretin, which is known to dramatically affect membrane electrical and elastic properties^33-37^. In our previous studies we already demonstrated that this strategy worked successfully with polyene antifungals, amphotericin B and nystatin, antifungal lipopeptide syringomycin E and antibacterial lipopeptide antibiotic polymyxin B^21,22,38,39,40^.

Figure 8a demonstrates the effects of 20 μM phloretin on the transmembrane current induced by MCF in the POPC:Erg (67:33 mol.%) membranes bathed in asymmetric salt solutions at 100 mV. Phloretin led to ten-fold increase in the steady-state MCF-induced transmembrane current. Phloretin did not alter the value of the *V*_*rev*_ of bilayers treated with MCF (Figure 8b). Moreover, step-like current fluctuations were observed in the presence of phloretin (Figure 8a, Figure 8b, insets) that should be assigned to functioning of MCF-produced ion-permeable pores. These data clearly demonstrated that MCF was able to potentiate pore-forming activity of MCF. Phloretin is known to decrease the membrane dipole potential of lipid bilayers^35,41,42^. The decrease in the dipole potential (with the hydrocarbon region being positive relative to the aqueous phase) is expected to diminish the electrostatic energy at the center of the pore for cations and increase for anions^43^. This should cause an increase in the steady-state transmembrane currents induced by cationic agents such as syringomycin E and polymyxin B^39,40^. Taking into account that molecules of MCF possess negative net charge, the increase in the number of MCF channels at the phloretin introduction was not a result of reduction of bilayer dipole potential. Considering that phloretin also decrease a lipid packing stress in the membrane^44^, we proposed that the increase in the number of MCF channels was due to a perturbation in the elastic properties of the membrane.

We also examined the potential antifungal effect of liposomal forms of MCF without and with the potentiator of its pore forming activity, phloretin. Clinical isolates of *Candida* species resistant to fluconazole were used, except for the *C. neoformans* strain (sensitive to fluconazole). Amphotericin B dissolved in DMSO was used as a positive control sample. Liposomes without MCF and phloretin and lipid vesicles with phloretin alone did not demonstrate any antifungal activity against test microorganisms at the concentrations used. Table 3 demonstrates that combination of MCF with phloretin resulted in decrease of antibiotic minimum inhibitory concentration against *Candida albicans* 604M, *Candida tropicalis* 56-05, and *Candida krusei* 72-05 compared to liposomal form of MCF without phloretin.

**Table 3.** Antifungal activity was assessed visually after 24/48 hours of incubation at 35°C. The minimum inhibitory concentration values corresponded to the minimum concentration at which there was no visible growth of the test microorganism.

|  | minimum inhibitory concentration (µg/mL) (24/48 hours) |  |  |
| --- | --- | --- | --- |
|  | **MCF** | **MCF with phloretin** | **amphotericin B** |
| *C. albicans* 604M | 0.06/0.06 | ≤0.03/ ≤0.03 | 1/1 |
| *C. tropicalis* 56-05 | 0.06/0.125 | ≤0.03/ ≤0.03 | 1/2 |
| *C. krusei* 72-05 | 0.25/0.25 | 0.06/0.06 | 2/2 |
| *C. glabrata* 61L | ≤0.03/≤0.03 | ≤0.03/≤0.03 | 1/2 |
| *Cryptococcus neoformans* n\ n | ≤0.03/0.06 | ≤0.03/≤0.03 | 1/1 |

We think that combinations of echinocandins with plant-derived small molecules that increase their pore-forming activity could facilitate the development of innovative forms of antibiotics that have two different mechanisms of action, both by inhibiting GS and forming pores in the membranes of pathogenic fungi.

## Conclusions

Our study provides an insight into the critical role of membrane activity of echinocandins. The dramatical changes in the membrane permeability at the adsorption of echinocandins and the ability of antibiotics to form ion-permeable pores in the model lipid membranes have been demonstrated. This might indicate an additional mechanism of antifungal action of these drugs. We have shown for the first time that the echinocandins (anidulafungin, caspofungin, and micafungin) affect membrane domain organization and the thermotropic characteristics of sterol-containing membranes, leading to the formation of quasi-gel lipid phase. Certainly, these effects might play a role in the (1,3) beta-glucan synthase functioning. We also discovered a way to potentiate the pore-forming activity of micafungin by combining it with the plant polyphenol phloretin, which resulted in a significant reduction in the minimum inhibitory concentration against flucanozole-resistant *Candida albicans, Candida tropicalis*, and *Candida krusei*. Detected membrane activity of echinocandins could promote the development of new therapeutic target agents to combat antibiotic resistance.

## References

(1) Rodrigues, M.L., & Nosanchuk, J.D. Fungal diseases as neglected pathogens: A wake-up call to public health officials. PLoS Negl Trop Dis. 14(2), e0007964 (2020). DOI: 10.1371/journal.pntd.0007964.

(2) Kainz, K., Bauer, M.A., Madeo, F., & Carmona-Gutierrez, D. Fungal infections in humans: the silent crisis. Microb Cell. 7(6), 143–145 (2020). DOI: 10.15698/mic2020.06.718.

(3) WHO fungal priority pathogens list to guide research, development and public health action. Geneva: World Health Organization. 2022. Licence: CC BY-NC-SA 3.0 IGO.

(4) Pappas, P.G., Kauffman, C.A., Andes, D.R., Clancy, C.J., Marr, K.A., Ostrosky-Zeichner, L., Reboli, A.C., Schuster, M.G., Vazquez, J.A., Walsh, T.J., Zaoutis, T.E., & Sobel, J.D. Clinical Practice Guideline for the Management of Candidiasis: 2016 Update by the Infectious Diseases Society of America. Clin Infect Dis. 62(4), e1–50 (2016). DOI: 10.1093/cid/civ933.

(5) Htet, L.L., Wang, L.N., & Liew, Y.X. Efficacy and safety of echinocandins versus triazoles or amphotericin B in the treatment of invasive fungal infections in paediatric patients: a systematic review. Singapore Med J. 2023. DOI: 10.4103/singaporemedj.SMJ-2021-173.

(6) Tóth, Z., Bozó, A., Kovács, R., Balogh, B., Balázs, B., Forgács, L., Kelentey, B., & Majoros, L. The In Vitro Activity of Fluconazole, Amphotericin B and Echinocandins Against Cyberlindnera fabianii Planktonic Cells and Biofilms. Mycopathologia, 188(1-2), 111–118 (2023). DOI: 10.1007/s11046-022-00688-9.

(7) Szymański, M., Chmielewska, S., Czyżewska, U., Malinowska, M., & Tylicki, A. Echinocandins - structure, mechanism of action and use in antifungal therapy. J Enzyme Inhib Med Chem. 37(1), 876–894 (2022). DOI: 10.1080/14756366.2022.2050224.

(8) Hu, X., Yang, P., Chai, C., Liu, J., Sun, H., Wu, Y., Zhang, M., Zhang, M.,Liu, X., & Yu H. Structural and mechanistic insights into fungal β-1,3-glucan synthase FKS1. Nature. 616(7955), 190–198 (2023). DOI: 10.1038/s41586-023-05856-5.

(9) Zhao, C.R., You, Z.L., Chen, D.D., Hang, J., Wang, Z.B., Ji, M., Wang, L.X., Zhao, P., Qiao, J., Yun, C.H., & Bai, L. Structure of a fungal 1,3-β-glucan synthase. Sci Adv. 9(37), eadh7820 (2023). DOI: 10.1126/sciadv.adh7820.

(10) Healey, K.R., Katiyar, S.K., Raj, S., & Edlind, T.D. CRS-MIS in Candida glabrata: sphingolipids modulate echinocandin-Fks interaction. Mol Microbiol. 86(2), 303–313 (2012). DOI: 10.1111/j.1365-2958.2012.08194.x.

(11) Satish, S., Jiménez-Ortigosa, C., Zhao, Y., Lee, M.H., Dolgov, E., Krüger, T.;, Park, S., Denning, D.W., Kniemeyer, O., Brakhage, A.A., & Perlin, D.S. Stress-Induced Changes in the Lipid Microenvironment of β-(1,3)-d-Glucan Synthase Cause Clinically Important Echinocandin Resistance in Aspergillus fumigatus. mBio, 10(3), e00779–19 (2019). DOI: 10.1128/mBio.00779-19.

(12) Zheng, L., Xu, Y., Wang, C., Yang, F., Dong, Y., & Guo, L. Susceptibility to caspofungin is regulated by temperature and is dependent on calcineurin in Candida albicans. Microbiol Spectr. 11(6), e0179023 (2023). DOI: 10.1128/spectrum.01790-23.

(13) Moreira-Walsh, B., Ragsdale, A., Lam, W., Upadhya, R., Xu, E., Lodge, J.K., & Donlin, M.J. Membrane Integrity Contributes to Resistance of Cryptococcus neoformans to the Cell Wall Inhibitor Caspofungin. mSphere. 7(4), e0013422 (2022). DOI: 10.1128/msphere.00134-22.

(14) Lockhart, S.R., Chowdhary, A., & Gold, J.A.W. The rapid emergence of antifungal-resistant human-pathogenic fungi. Nat Rev Microbiol. 21(12), 818–832 (2023). DOI: 10.1038/s41579-023-00960-9.

(15) Rhodes, J., & Fisher, M.C. Global epidemiology of emerging Candida auris. Curr Opin Microbiol. 52, 84–89 (2019). DOI: 10.1016/j.mib.2019.05.008.

(16) Lockhart, S.R., Etienne, K.A., Vallabhaneni, S., Farooqi, J., Chowdhary, A., Govender, N.P., Colombo, A.L., Calvo, B., Cuomo, C.A., Desjardins, C.A., Berkow, E.L., Castanheira, M., Magobo, R.E., Jabeen, K., Asghar, R.J., Meis, J.F., Jackson, B., Chiller, T., & Litvintseva, A.P. Simultaneous Emergence of Multidrug-Resistant Candida auris on 3 Continents Confirmed by Whole-Genome Sequencing and Epidemiological Analyses. Clin Infect Dis. 64(2), 134–140 (2017). DOI: 10.1093/cid/ciw691.

(17) Tripathi, S.K., Feng, Q., Liu, L., Levin, D.E., Roy, K.K., Doerksen, R.J., Baerson, S.R., Shi, X., Pan, X., Xu, W.H., Li, X.C., Clark, A.M., & Agarwal, A.K. Puupehenone, a Marine-Sponge-Derived Sesquiterpene Quinone, Potentiates the Antifungal Drug Caspofungin by Disrupting Hsp90 Activity and the Cell Wall Integrity Pathway. mSphere. 5(1), e00818–19 (2020). DOI: 10.1128/mSphere.00818-19.

(18) Sun, Q., Xiong, K., Yuan, Y., Yu, J., Yang, L., Shen, C., Su, C., Lu, Y. Inhibiting Fungal Echinocandin Resistance by Small-Molecule Disruption of Geranylgeranyltransferase Type I Activity. Antimicrob Agents Chemother. 64(2), e02046–19 (2020). DOI: 10.1128/AAC.02046-19.

(19) Liu, N., Zhang, N., Zhang, S., Zhang, L., Liu, Q. Phloretin inhibited the pathogenicity and virulence factors against Candida albicans. Bioengineered. 12(1), 2420–2431 (2021). DOI: 10.1080/21655979.2021.1933824.

(20) Cheon, D., Kim, J., Jeon, D., Shin, H.C., & Kim, Y. Target Proteins of Phloretin for Its Anti-Inflammatory and Antibacterial Activities Against Propionibacterium acnes-Induced Skin Infection. Molecules. 24(7), 1319 (2019). DOI: 10.3390/molecules24071319.

(21) Chulkov, E.G., Schagina, L.V., & Ostroumova, O.S. Membrane dipole modifiers modulate single-length nystatin channels via reducing elastic stress in the vicinity of the lipid mouth of a pore. Biochim Biophys Acta. 1848(1PtA), 192–199 (2015). DOI: 10.1016/j.bbamem.2014.09.004.

(22) Efimova, S.S., Malykhina, A.I., & Ostroumova, O.S. Triggering the Amphotericin B Pore-Forming Activity by Phytochemicals. Membranes. 13(7), 670 (2023). DOI: 10.3390/membranes13070670.

(23) Pierce, C.G., Srinivasan, A., Uppuluri, P., Ramasubramanian, A.K., & López-Ribot, J.L. Antifungal therapy with an emphasis on biofilms. Curr Opin Pharmacol. 13(5), 726–730 (2013). DOI: 10.1016/j.coph.2013.08.008.

(24) Crump, J.A., & Collignon, P.J. Intravascular catheter-associated infections. Eur J Clin Microbiol Infect Dis. 19(1), :1-8 (2000). DOI: 10.1007/s100960050001.

(25) Hassanpour, P., Hamishehkar, H., Bahari Baroughi, B., Baradaran, B., Sandoghchian Shotorbani, S., Mohammadi, M., Shomali, N., Aghebati-Maleki, L., & Nami, S. Antifungal Effects of Voriconazole-Loaded Nano-Liposome on Fluconazole-Resistant Clinical Isolates of Candida albicans, Biological Activity and ERG11, CDR1, and CDR2 Gene Expression. Assay Drug Dev Technol. 19(7), 453–462 (2021). DOI: 10.1089/adt.2020.1057.

(26) Maertens, J., Pagano, L., Azoulay, E., & Warris, A. Liposomal amphotericin B-the present. J Antimicrob Chemother. 77(Suppl_2), ii11–ii20 (2022). DOI: 10.1093/jac/dkac352.

(27) Vera-González, N., Bailey-Hytholt, C.M., Langlois, L., de Camargo Ribeiro, F., de Souza Santos, E.L., Junqueira, J.C., & Shukla, A. Anidulafungin liposome nanoparticles exhibit antifungal activity against planktonic and biofilm Candida albicans. J Biomed Mater Res A. 108(11), 2263–2276 (2020). DOI: 10.1002/jbm.a.36984.

(28) Singh, A., Yadav, V., & Prasad, R. Comparative lipidomics in clinical isolates of Candida albicans reveal crosstalk between mitochondria, cell wall integrity and azole resistance. PLoS One. 7(6), e39812 (2012). DOI: 10.1371/journal.pone.0039812.

(29) Löffler, J., Einsele, H., Hebart, H., Schumacher, U., Hrastnik, C., & Daum, G. Phospholipid and sterol analysis of plasma membranes of azole-resistant Candida albicans strains. FEMS Microbiol Lett. 185(1), 59–63 (2000). DOI: 10.1111/j.1574-6968.2000.tb09040.x.

(30) van Meer, G., & de Kroon, A.I. Lipid map of the mammalian cell. J Cell Sci. 124(Pt 1), 5–8 (2011). DOI: 10.1242/jcs.071233.

(31) Luchini, A., & Vitiello, G. Mimicking the Mammalian Plasma Membrane: An Overview of Lipid Membrane Models for Biophysical Studies. Biomimetics. 6(1), 3 (2020). DOI: 10.3390/biomimetics6010003.

(32) Fujie, Akihiko. “Discovery of micafungin (FK463): A novel antifungal drug derived from a natural product lead” Pure and Applied Chemistry, vol. 79, no. 4, 2007, pp. 603–614. DOI: 10.1351/pac200779040603.

(33) Cseh, R., & Benz, R. Interaction of phloretin with lipid monolayers: relationship between structural changes and dipole potential change. Biophys J. 77(3), 1477–1488 (1999). DOI: 10.1016/S0006-3495(99)76995-X.

(34) Ostroumova, O.S., Efimova, S.S., & Schagina, L.V. Phloretin-induced reduction in dipole potential of sterol-containing bilayers. J Membr Biol. 246(12), 985–991 (2013). DOI: 10.1007/s00232-013-9603-2.

(35) Efimova, S.S., & Ostroumova, O.S. Effect of dipole modifiers on the magnitude of the dipole potential of sterol-containing bilayers. Langmuir. 28(26), 9908–9914 (2012). DOI: 10.1021/la301653s.

(36) Ostroumova, O.S., Chulkov, E.G., Stepanenko, O.V., & Schagina, L.V. Effect of flavonoids on the phase separation in giant unilamellar vesicles formed from binary lipid mixtures. Chem Phys Lipids. 178, 77–83 (2014). DOI: 10.1016/j.chemphyslip.2013.12.005.

(37) Efimova, S.S., Malev, V.V., & Ostroumova, O.S. Effects of Dipole Potential Modifiers on Heterogenic Lipid Bilayers. J Membr Biol. 249(1-2), 97–106 (2016). DOI: 10.1007/s00232-015-9852-3.

(38) Chulkov, E.G., & Ostroumova, O.S. Phloretin modulates the rate of channel formation by polyenes. Biochim Biophys Acta. 1858(2), 289–294 (2016). DOI: 10.1016/j.bbamem.2015.12.004.

(39) Ostroumova, O.S., Shchagina, L.V., & Malev, V.V. The effect of dipole potential of lipid bilayers on the properties of ion channels formed by cyclic lipodepsipeptide syringomycin E. Biochem. (Moscow) Suppl. Ser. A: Membr. Cell Biol. 2, 259–270 (2008). DOI: 10.1134/s1990747808030100.

(40) Zakharova, A.A., Efimova, S.S., & Ostroumova, O.S. Lipid Microenvironment Modulates the Pore-Forming Ability of Polymyxin B. Antibiotics. 11(10), 1445 (2022). DOI: 10.3390/antibiotics11101445.

(41) Franklin, J.C., & Cafiso, D.S. Internal electrostatic potentials in bilayers: measuring and controlling dipole potentials in lipid vesicles. Biophys J. 65(1), 289–299 (1993). DOI: 10.1016/S0006-3495(93)81051-8.

(42) Pohl, P., Rokitskaya, T.I., Pohl, E.E., & Saparov, S.M. Permeation of phloretin across bilayer lipid membranes monitored by dipole potential and microelectrode measurements. Biochim Biophys Acta. 1323(2), 163–72 (1997). DOI: 10.1016/s0005-2736(96)00185-x.

(43) Andersen, O.S., Finkelstein, A., Katz, I., & Cass, A. Effect of phloretin on the permeability of thin lipid membranes. J Gen Physiol. 67(6), 749–771 (1976). DOI: 10.1085/jgp.67.6.749.

(44) Zlodeeva, P.D., Shekunov, E.V., Ostroumova, O.S., & Efimova, S.S. The Degree of Hydroxylation of Phenolic Rings Determines the Ability of Flavonoids and Stilbenes to Inhibit Calcium-Mediated Membrane Fusion. Nutrients. 15(5), 1121 (2023). DOI: 10.3390/nu15051121.

