## Supplementary material for "Echinocandins have an alternative mode of action on biomimetic membranes that is not directly related to the functioning of (1,3) beta-glucan synthase": Materials and methods, Supporting Figures and Tables

All chemicals were of reagent grade. Synthetic 1-palmitoyl-2-oleyl-*sn*-glycero-3-phoshocholine (POPC), 1,2-dipalmitoyl-*sn*-glycero-3-phosphocholine (DPPC), 1,2-dioleoyl-sn-glycero-3-phosphocholine (DOPC), 1,2-dipalmitoyl-*sn*-glycero-3-phosphoethanolamine-N-(lissamine rhodamine B sulfonyl) (Rh-DPPE), and cholesterol (Chol) were obtained from Avanti Polar Lipids, Inc (Pelham, AL). Caspofungin (CSF) (≥97% (HPLC)), anidulafungin (ANF) (≥97% (HPLC)), micafungin (MCF) (≥97% (HPLC)), ergosterol (Erg), calcein, dimethylsulfoxide (DMSO), sephadex G-50, triton X-100, sorbitol, EDTA, NaCl, HEPES, KOH and NaOH were purchased from Aldrich Company Ltd. (Gillingham, United Kingdom). Stock solutions of echinocandins were prepared with DMSO. Distilled water was used.

**Calcein release from lipid unilamellar vesicles**

The fluorescence of calcein that leaked from lipid unilamellar vesicles (LUVs) was used to monitor the membrane permeabilization induced by echinocandins. LUVs were prepared from POPC:Erg (67:33 mol.%), POPC:Chol (67:33 mol.%) or pure POPC by extrusion using an Avanti Polar Lipid mini-extruder (Pelham, AL). The lipid stock in chloroform was dried under a gentle stream of nitrogen. A dry lipid film was hydrated by a calcein-containing buffer (35 mM calcein, 10 mM HEPES-NaOH, pH 7.4). The suspension was subjected to five freeze-thaw cycles and passed through a 100-nm nucleopore polycarbonate membrane 13 times. Calcein that was not entrapped in vesicles was removed by gel filtration in a sephadex G-50 column to replace the buffer outside the liposomes with a calcein-free solution (0.15 M NaCl, 1 mM EDTA, 10 mM HEPES-NaOH, pH 7.4). The concentration of the obtained LUV suspension was 3 mM. The calcein in vesicles fluoresced very poorly due to strong self-quenching at millimolar concentrations, while the fluorescence of the disengaged calcein in the surrounding solution correlated with membrane permeabilization in the absence and presence of echinocandins.

Echinocandins were added to calcein-loaded liposomes from the stock solution (4 mM in DMSO). Time-dependent calcein fluorescence de-quenching induced by 50 μM of echinocandins was measured over 50 min, at least three repeats for each lipid/echinocandin combination.

The degree of calcein release was determined at 25 ± 2°C using a Fluorat-02-Panorama spectrofluorimeter (Lumex, Saint-Petersburg, Russia). A 10-mm quartz cuvette was used to measure calcein release from liposomes after the addition of echinocandins. The excitation wavelength was 490 nm, and the emission wavelength was 520 nm. Addition of triton X-100 from a 10 mM water solution to a final concentration of 1% to each sample led to complete disruption of LUVs, and the intensity of fluorescence after releasing the total amount of calcein from liposomes was measured.

The relative intensity of calcein fluorescence (*IF*, %) was used to describe the dependence of permeabilization of the liposomes on the type of echinocandin and membrane composition. *IF* was calculated using the following formula:

$$IF = \frac{I - I_0}{\frac{Imax}{0.9} - I_0} \cdot 100\%, \quad (1)$$

where $I$ and $I_0$ are the calcein fluorescence intensities in the sample in the presence and in the absence of an echinocandin, respectively, and $I_{max}$ is the maximal fluorescence of the sample after lysis of liposomes by triton X-100. A factor of 0.9 was introduced to calculate the dilution of the sample by detergent.

The kinetics of calcein release was described by a one-exponential function with time constant, τ. The values of statistical criterion $R^2$ were in the range of 0.8-0.9. The values of $IF_{\max}$ and τ were averaged from 3 to 5 independent experiments and are presented as mean ± standard error ($p \leq 0.05$).

**Permeability of planar lipid bilayer**

Virtually solvent-free bilayers were prepared according to a monolayer-opposition technique[1] on a 50 μm-diameter aperture in a 10 μm-thick Teflon film, which separates the Teflon chamber into two distinct compartments (*cis*- and *trans*-) compartments. The aperture was pretreated with hexadecane. The planar lipid membranes were composed of mixtures of 67 mol.% POPC and 33 mol.% sterol (Erg or Chol) and bathed in asymmetric salt solutions of 0.025 M NaCl

(10 mM HEPES, pH 7.4 (*cis*-side)) and 0.15 M NaCl (10 mM HEPES, pH 7.4 (*trans*-side)). After the membrane was completely formed and stabilized, echinocandins were added to the *cis*-compartment from a 4 mM in DMSO up to a final concentration from 0.5 to 5 µM. Solvent alone did not affect membrane stability and permeability. The measurements of the transmembrane current (*I*) and the applying the transmembrane voltage (*V*) were produced using Ag/AgCl electrodes with 2 M KCl/1.5% agarose bridges. At positive voltage *cis*-compartment was positive with respect to the *trans*-compartment. All experiments were performed at room temperature (25 ± 2°C).

The Axopatch 200B amplifier (Molecular Devices, LLC, Orleans Drive, Sunnyvale, CA, USA) in the voltage-clamp mode was used to measure the current. The signals were digitized by Digidata 1440A (Molecular Devices, LLC, Orleans Drive, Sunnyvale, CA, US) at a 5 kHz sampling frequency using 1 kHz low-pass filtering. The data analysis was performed using pClamp 10 (Molecular Devices, LLC, Orleans Drive, Sunnyvale, CA, USA) at filtering by an 8-pole Bessel 100 kHz and Origin 8.0 (OriginLab Corporation, Northampton, MA, USA).

Transport numbers of cations ($t^+$) and anions ($t^- = 1 - t^+$) were determined as[2]:

$$V^{rev} = \frac{kT}{e}(1 - 2t^+)\, ln\left(\frac{C_1}{C_2}\right), \tag{2}$$

where $V^{rev}$ is the reversal potential (the voltage at which transmembrane current is equal to 0); *k, T, e* have their usual meanings; $C_1$ and $C_2$ are the concentrations of NaCl solutions in the *cis*- and *trans*- compartments, respectively.

To enhance MCF pore-forming activity phloretin was added at the both-side of POPC:Erg (67:33 mol.%) membrane treated with antibiotic up to 20 µM. Four independent experiments were carried out. Phloretin was not able to increase the ionic permeability of lipid bilayers alone.

**Differential scanning microcalorimetry**

Giant unilamellar vesicles were prepared from pure DPPC and mixture of DPPC:Chol (85:15 mol.%) and DPPC:Erg (85:15 mol.%) using the electroformation method with Vesicle Pre Pro® (Nanion Technologies, Munich, Germany) (standard protocol, 3 V, 10 Hz, 58 min, 55°C). The obtained liposome suspension contained 5 mM lipid and was buffered by 5 mM HEPES-KOH at pH 7.4. Echinocandins, ANF, CSF and MCF, from mM stock solutions in DMSO were added to aliquots to the lipid:echinocandin molar ratio of 50:1, 25:1 and 10:1. Then suspension was heated and cooled at constant rates of 0.2 and 0.3°C/min, respectively, at a µDSC 7EVO microcalorimeter (Setaram, Caluire-et-Cuire, France). The reversibility of the thermal transitions was assessed by reheating the sample immediately after the cooling step from the previous scan.

The lipid thermograms were characterized by the temperature of pre-transition attributed to the mobility of the choline polar head (only in the case of DPPC vesicles), the mean melting temperature (the temperature at which excess heat capacity reaches a maximum, $T_m$) and the enthalpy of the main phase transition (an area under the main peak, $\Delta H_{cal}$). The sharpness of the gel-to-liquid-crystalline phase transition was expressed as the temperature difference between the upper (onset) and lower (completion) boundary of the main phase transition, $\Delta T_b$.

The deconvolution analysis of main peak in the presence of echinocandins was performed using Calisto software. The separation of multiple overlapped peaks was based on the application of Gaussian and/or Fraser-Suzuki (asymmetric) type signals. The melting temperature of each *i*-peak component, $T_{m\_i}$, and the percentage contribution of *i*-peak component to the total area, $\boldsymbol{p_i} = \frac{\boldsymbol{\Delta H_i/\Delta H_{cal}}}{\sum_i \boldsymbol{\Delta H_i/\Delta H_{cal}}}$, were determined. The fitting of calculated signal to the experimental data was performed using a non-linear optimization (Marquardt).

To investigate the dependences of thermotropic behavior of DPPC, DPPC:Chol (85:15 mol.%) and DPPC:Erg (85:15 mol.%) on echinocandin type mean melting temperature, $T_{m\_mean}$, was determined as:

$$T_{m\_mean} = \frac{\sum_i T_{m\_i} \cdot p_i}{\sum_i p_i}. \tag{3}$$

At least two independent experiments were done for each lipid composition/echinocandin combination to ensure the reproducibility of the results obtained.

**Confocal microscopy of giant unilamellar vesicles**

Giant unilamellar vesicles (GUVs) were formed from DOPC:Chol (80:20 mol.%) mixture by the electroformation method as described above (standard protocol, 3 V, 10 Hz, 58 min, 45°C). Lipid stock solutions was prepared in chloroform. Labeling was carried out by addition of the fluorescent lipid probe, Rh-DPPE (1 mol.%). The resulting aqueous liposome suspension containing 0.8 mM lipid and 1.5 M sorbitol was divided into 30 µl aliquots. The control liposome samples did not contain echinocandins. ANF and MCF were introduced into liposome suspension from DMSO stock solutions up to 50 µM. CSF was tested at 10 µM, while it had detergent effect[3]. The liposome suspension with echinocandins was allowed to equilibrate for 30 min at room temperature (25 ± 2°C). The sample was observed as a standard microscopy preparation. 10 µl of the resulting liposome suspension was placed on a standard microscope slide and covered by a coverslip. GUVs were imaged through an oil immersion objective (65×/1.4HCX PL) using an Olympus (Hamburg, Germany). Temperature during observation was controlled by the air heating/cooling in the thermally insulated camera. Rh-DPPE was excited at wavelengths of 543 nm (helium–neon laser).

Rh-DPPE clearly favors liquid disordered phase and it is excluded from ordered phases[4]. The existence of surface tension forces, tending to shorten the length of the boundary between the ordered and disordered phases, determines a circular shape of liquid ordered regions that appear as uncolored by Rh-DPPE round domains. Uncolored domains of complex shape should be referred to gel lipid phase. Predominantly, DOPC:Chol (80:20 mol.%) GUVs were visually homogeneous with liquid disordered lipid phase. Few vesicles might contain ordered lipid phase which is described for DOPC:Chol (70:30 mol.%)[5]. Numbers of GUVs within single field of view without and with visible phase separation of various types were counted. Several neighboring fields of view were analyzed.

Part of vesicles with different phase separation at each tested system was calculated as the ratio of particular phase GUVs to the total number of GUVs. Experiments were carried out in 8 replicates with mean number of liposomes per condition – 125. Difference between groups was assessed by one-way ANOVA test with significance level $p \leq 0.05$.

**Molecular dynamics simulation**

Three model membranes were assembled in CHARMM-GUI Membrane Builder[6] and contained following number of lipid molecules: (1) 50 POPC; (2) 40 POPC, 20 Chol; (3) 40 POPC, 20 Erg. Water solution was ionized with NaCl to concentration 0.15 M. Echinocandins topology parameters were generated using CGenFF[7]. One molecule of ANF, CSF or MCF was placed at 3 nm above membrane center. The size of the simulation box was approximately 4x4x13 nm.

GROMACS 2023.2[8] was used to perform MD simulation with CHARMM36m all-atom force field[9]. Energy minimization was carried out by the steepest descent algorithm. A six-step equilibration process was performed by gradually turning off the position restraints on lipid molecules. All simulations were conducted at constant temperature 25°C (298.15 K) and pressure 1 bar using the Nosé-Hoover and semi-isotropic pressure coupling approach with C-rescale barostat[10,11]. The time constant of coupling for temperature and pressure was 1 and 5 ps, respectively. The Particle Mesh Ewald method was employed to treat long-range electrostatic interaction with a short-range cut-off 1.2 nm[12], while the shifted Lennard-Jones potential algorithm was used to calculate the Van der Waals interactions with a general cut-off of 1.2 nm and a shifting cut-off of 1.0 nm. The trajectory in production simulations was recorded every 10 ps, in steered simulations and umbrella sampling – every 1 ps. Production simulations were performed for 100 ns. Area per lipid headgroups (APL) in POPC membrane was 63.1Å$^2$ which is in good agreement with experimental (62.7 Å$^2$ at 20°C, 64.3 Å$^2$ at 30°C[13]) and MD data (63.5 Å$^2$ at 303.15 K[14]).

Steered MD simulations were performed in order to enhance the penetration of echinocandins through membranes. This method is the most extensively used to characterize molecule-membrane interaction and obtain related free energy profiles[15-17]. In this study, a harmonic potential with a force constant of 500 kJ/mol/nm$^2$ was applied between the center of mass of the echinocandin molecule and the membrane along the Z-axis. The frame, where echinocandin was 3 nm above the membrane center, was chosen from production run and was then used as the starting point for 50 ns pulling which enabled the molecule to penetrate the lipid bilayer smoothly. Successive umbrella sampling windows were conducted for 50 ns at distances from 0 to 3.0 nm away from membrane center along the Z-axis at every 0.2 nm, resulting in 16 simulation windows. The weighted histogram analysis method (WHAM) was then applied to the resulting data to obtain the potential of mean force (PMF)[18]. Standard error was calculated by the Bootstrap method[19].

Echinocandins influence on membrane properties was assessed in separate simulations. One molecule of the drug was embedded in model membrane in such a way that its head was located in lipid headgroup region while hydrophobic part of the molecule was in lipid tail core. The size of the simulation box was approximately 6x6x8 nm. All other settings were the same. Production simulations were performed for 100 ns. APL was calculated by MEMBPLUGIN[20].

**Antifungal activity**

*Preparation of the of liposomal forms*

The technique by[21] with modifications for the formation of the liposomes treated with MCF was used. Lipid mixture (67 mol.% POPC and 33 mol.% Chol) without or with MCF and/or phloretin were suspended in a mixture of chloroform (67 vol.%) and methanol (33 vol.%). The equimolar contents of lipid, MCF and phloretin were used. The resulting solution was evaporated in a vacuum rotary evaporator at room temperature for 120 min. Further, the lipid film was dispersed in a buffer (0.15 M NaCl, 10 mM HEPES, 1 mM EDTA, pH 7.4) and was exposed to ultrasound for 10 min. The concentration of MCF in samples used for antifungal tests was assessed by measuring UV-spectra basing on the method proposed by[22]. Absorbance spectroscopy was performed using a spectrofluorimeter "Fluorat-02-Panorama" (Lumex, Saint-Petersburg, Russia). All samples were scanned from 240 to 340 nm with a 1 nm step, the concentration MCF was determined by absorbance at the spectra maximum.

*Test microorganisms*

Clinical isolates of fluconazole-resistant *Candida albicans* 604M, *Candida tropicalis* 56-05, *Candida krusei* 72-05*, Candida glabrata* 61L, and fluconazole-sensitive *Candida neoformans* strain for the screening of antifungal compounds were kindly provided by A.B. Kulko (Moscow Government Health Department Scientific and Clinical Antituberculosis Center, Moscow).

*Sample preparation*

Samples were diluted in RPMI1640 medium to a concentration of 8 μg/mL, according to the content of the active component. Amphotericin B, which was used as a control compound, was dissolved in DMSO to a concentration of 10,000 μg/mL in accordance with the percentage of the main substance (95%) and further diluted in the nutrient medium. The working concentration range of tested samples was 4-0.03 μg/mL. The range of working concentrations of amphotericin B was 32-0.25 μg/mL.

*Assay setup*

Antifungal activity was studied by the serial microdilution method in the nutrient broth according to EUCAST Definitive Document E.DEF 7.3.2 (**EUCAST Definitive document E.DEF 7.3.2 (April 2020) Method for the determination of broth dilution minimum inhibitory concentrations of antifungal agents for yeasts**). The nutrient medium RPMI 1640 (with L-glutamine and pH indicator but without bicarbonate) was prepared according to the normative documentation from dry powder (SIGMA Lot SLBZ6264, St. Loius, USA) with the addition of glucose at a final concentration of 2% and MOPS (3-(N-morpholino)propanesulfonic acid, PanEco, Russia) to a final concentration of 0.165 mol/L. The pH was adjusted to 7.0 with 1 M/L sodium hydroxide.

For each Candida culture from single colonies grown on 48-hour Sabouraud agar, were suspended in PBS solution, and the cell inoculum titer was determined according to the McFarland standard (equal to $5\times10^6$ CFU $ml^{-1}$) of the microbial suspension using a Den-1B densitometer (suspension turbidity detector) (Latvia). After a series of dilutions of the tested samples in 96-well plates (Medpolymer, Russia), inoculum was added to a final titer of about $2\times10^5$ CFU/mL The plates were incubated without shaking at 35±2°C under aerobic conditions.

Antifungal activity was assessed visually after 24 and 48 hours. The MIC values corresponded to the minimum concentration at which there was no visible growth of the test microorganism.

### Supporting Figures and Tables

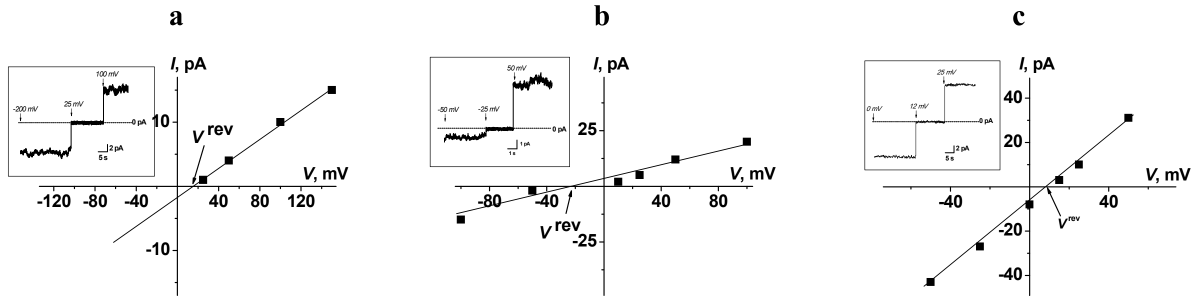

**Figure S1.** Cation/anion selectivity of transmembrane pores induced by ANF (a), CSF (b) and MCF (c). I–V curves of modified with sufficient amount of echinocandins (cis-side only) to induce a bilayer permeability. The membranes were composed of POPC:Chol (67:33 mol.%) and bathed in asymmetric salt solutions of 0.025 M NaCl (10 mM HEPES, pH 7.4 (cis-side)) and 0.15 M NaCl (10 mM HEPES, pH 7.4 (trans-side)). Arrows indicate the reversal potential ($V_{rev}$). Inset: Current fluctuations of the ANF (a), CSF (b) and MCF (C) pores at different transmembrane voltages in asymmetric solutions.

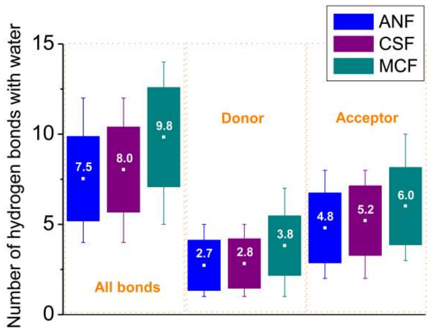

**Figure S2.** Average number of hydrogen bonds formed between ANF (blue squares), CSF (violet squares) or MCF (green squares) molecule and water during 100 ns molecular dynamic simulation in water box (0.15 M NaCl). Bonds where echinocandin acts as donor or acceptor showed separately.

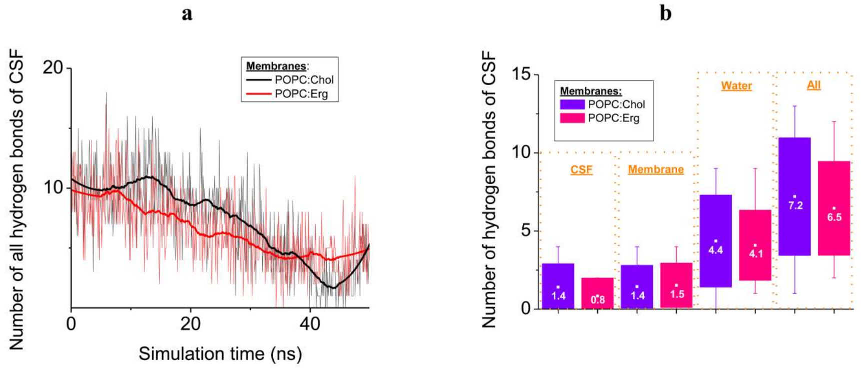

**Figure S3.** Analysis of hydrogen bonds formed by CSF in POPC:Chol (67:33 mol.%) and POPC:Erg (67:33 mol.%) membranes during steered molecular dynamic simulation, the molecule reach membrane center on approximately 35 ns. (a) Number of hydrogen bonds in time, bold lines represent averaging of the graphs. (b) Mean number of bonds formed between CSF and different components (CSF itself, membrane, water) and overall.

**Table S1.** The main peak decomposition/deconvolution analysis in the presence of echinocandins.

| | | | **DPPC** | | **DPPC:Chol (85:15 mol.%)** | | **DPPC:Erg (85:15 mol.%)** | |
|---|---|---|---|---|---|---|---|---|
| ***agent*** | ***ratio*** | ***peak*** | $T_{m\_i}$, °C | $\frac{\Delta H_i/\Delta H_{cal}}{\sum_i \Delta H_i/\Delta H_{cal}}$, % | $T_{m\_i}$, °C | $\frac{\Delta H_i/\Delta H_{cal}}{\sum_i \Delta H_i/\Delta H_{cal}}$, % | $T_{m\_i}$, °C | $\frac{\Delta H_i/\Delta H_{cal}}{\sum_i \Delta H_i/\Delta H_{cal}}$, % |
| ***ANF*** | – | *control* | 41.6 | 100 | 40.3 | 100 | 40.5 | 100 |
| | 50:1 | #1 | 41.6 | 80 | 40.0 | 100 | 40.4 | 74 |
| | | #2 | 41.0 | 20 | – | – | 39.6 | 26 |
| | 25:1 | #1 | 41.6 | 63 | 39.8 | 58 | 40.2 | 62 |
| | | #2 | 40.8 | 37 | 39.0 | 42 | 39.4 | 38 |
| | 10:1 | #1 | 41.9 | 58 | 39.7 | 44 | 40.0 | 59 |
| | | #2 | 41.3 | 31 | 38.9 | 56 | 38.1 | 41 |
| | | #3 | 39.7 | 11 | – | – | – | – |
| ***CSF*** | – | *control* | 41.5 | 100 | 40.8 | 100 | 40.4 | 100 |
| | 50:1 | #1 | 41.5 | 100 | 40.8 | 100 | 40.4 | 100 |
| | 25:1 | #1 | 41.6 | 100 | 40.9 | 100 | 40.4 | 100 |
| | 10:1 | #1 | 42.0 | 38 | 40.9 | 100 | 40.4 | 100 |
| | | #2 | 41.6 | 62 | – | – | – | – |
| ***MCF*** | – | *control* | 41.4 | 100 | 40.8 | 100 | 40.5 | 100 |
| | 50:1 | #1 | 41.5 | 54 | 41.0 | 24 | 40.2 | 100 |
| | | #2 | 40.8 | 46 | 40.4 | 76 | – | – |
| | 25:1 | #1 | 41.5 | 25 | 41.0 | 25 | 40.7 | 16 |
| | | #2 | 40.6 | 75 | 40.1 | 75 | 40.0 | 84 |
| | 10:1 | #1 | 42.0 | 24 | 40.7 | 36 | 41.1 | 24 |
| | | #2 | 40.3 | 76 | 39.7 | 64 | 39.7 | 76 |

**Supplemental Literature**

(1) Montal, M., & Mueller, P. Formation of bimolecular membranes from lipid monolayers and a study of their electrical properties. *Proc Natl Acad Sci USA*, **69**, 3561-3566 (1972). DOI: 10.1073/pnas.69.12.3561.

(2) Morf, W. Calculation of liquid-junction potentials and membrane potentials on the basis of the Planck theory. *Analytical Chemistry*, **49**(6), 810-813 (1977). DOI: 10.1021/ac50014a035.

(3) Sumiyoshi, M., Miyazaki, T., Makau, J.N., Mizuta, S., Tanaka, Y., Ishikawa, T., Makimura, K., Hirayama, T., Takazono, T., Saijo, T., Yamaguchi, H., Shimamura, S., Yamamoto, K., Imamura, Y., Sakamoto, N., Obase, Y., Izumikawa, K., Yanagihara, K., Kohno, S., & Mukae, H. Novel and potent antimicrobial effects of caspofungin on drug-resistant Candida and bacteria. *Sci Rep.*, **10**(1), 17745 (2020). DOI: 10.1038/s41598-020-74749-8.

(4) Juhasz, J., Davis, J.H., & Sharom, F.J. Fluorescent probe partitioning in giant unilamellar vesicles of 'lipid raft' mixtures. *Biochem J*, **430**(3), 415-423 (2010). DOI: 10.1042/BJ20100516.

(5) Suga, K., & Umakoshi, H. Detection of nanosized ordered domains in DOPC/DPPC and DOPC/Ch binary lipid mixture systems of large unilamellar vesicles using a TEMPO quenching method. *Langmuir*, **29**(15), 4830-4838 (2013). DOI: 10.1021/la304768f.

(6) Jo, S., Lim, J.B., Klauda, J.B., & Im, W. CHARMM-GUI Membrane Builder for mixed bilayers and its application to yeast membranes. *Biophys J*, **97**(1), 50-58 (2009). DOI: 10.1016/j.bpj.2009.04.013.

(7) Vanommeslaeghe, K., & MacKerell, A.D. Automation of the CHARMM General Force Field (CGenFF) I: bond perception and atom typing. *J Chem Inf Model*, **52**(12), 3144-3154 (2012). DOI: 10.1021/ci300363c.

(8) Abraham, M.J., Murtola, T., Schulz, R., Páll, S., Smith, J.C., Hess, B., & Lindahl, E. GROMACS: High performance molecular simulations through multi-level parallelism from laptops to supercomputers. *SoftwareX*, **(1-2)**, 19-25 (2015). DOI: 10.1016/j.softx.2015.06.001.

(9) Lee, J., Cheng, X., Swails, J.M., Yeom, M.S., Eastman, P.K., Lemkul, J.A., Wei, S., Buckner, J., Jeong, J.C., Qi, Y., Jo, S., Pande, V.S., Case, D.A., & Brooks, C.L. 3rd; MacKerell, A.D.Jr.; Klauda, J.B.; Im, W. CHARMM-GUI Input Generator for NAMD, GROMACS, AMBER, OpenMM, and CHARMM/OpenMM Simulations Using the CHARMM36 Additive Force Field. *J Chem Theory Comput*, **12**(1), 405-413 (2016). DOI: 10.1021/acs.jctc.5b00935.

(10) Martyna, G.J., Klein, M.L., & Tuckerman, M. Nosé–Hoover chains: The canonical ensemble via continuous dynamics. *J. Chem. Phys*, **97**(4), 2635–2643 (1992). DOI: 10.1063/1.463940.

(11) Bernetti, M., & Bussi, G. Pressure control using stochastic cell rescaling. *J Chem Phys*, **153**(11), 114107 (2020). DOI: 10.1063/5.0020514.

(12) Darden, T., York, D., & Pedersen, L. Particle mesh Ewald: An N·log(N) method for Ewald sums in large systems. *J. Chem. Phys*, **98**(12), 10089–10092 (1993). DOI: 10.1063/1.464397.

(13) Kučerka, N., Nieh, M.P., & Katsaras, J. Fluid phase lipid areas and bilayer thicknesses of commonly used phosphatidylcholines as a function of temperature. *Biochim Biophys Acta*, **1808**(11), 2761-2771 (2011). DOI: 10.1016/j.bbamem.2011.07.022.

(14) Saito, H., Morishita, T., Mizukami, T., Nishiyama, K., Kawaguchi, K., & Nagao, H. Molecular dynamics study of binary POPC bilayers: molecular condensing effects on membrane structure and dynamics. *Phys.: Conf. Ser*, **1136**, 012022 (2018). DOI: 10.1088/1742-6596/1136/1/012022.

(15) Lee, B.L., Kuczera, K., Middaugh, C.R., & Jas, G.S. Permeation of the three aromatic dipeptides through lipid bilayers: Experimental and computational study. *J Chem Phys*, **144**(24), 245103 (2016). DOI: 10.1063/1.4954241.

(16) Jiang, X., Yang, K., Yuan, B., Han, M., Zhu, Y., Roberts, K.D., Patil, N.A., Li, J., Gong, B., Hancock, R.E.W., Velkov, T., Schreiber, F., Wang, L., & Li, J. Molecular dynamics simulations informed by membrane lipidomics reveal the structure-interaction relationship of polymyxins with the lipid A-based outer membrane of Acinetobacter baumannii. *J Antimicrob Chemother*, **75**(12), 3534-3543 (2020). DOI: 10.1093/jac/dkaa376.

(17) Tran, D.P., Tada, S., Yumoto, A., Kitao, A., Ito, Y., Uzawa, T., & Tsuda, K. Using molecular dynamics simulations to prioritize and understand AI-generated cell penetrating peptides. *Sci Rep*, **11**(1), 10630 (2021). DOI: 10.1038/s41598-021-90245-z.

(18) Roux, B. The calculation of the potential of mean force using computer simulations. *Comp Phys Communication*, **91**(1–3), 275-282 (1995). DOI: 10.1016/0010-4655(95)00053-I.

(19) Hub, J.S., de Groot, B.L., & van der Spoel, D. g_wham—A Free Weighted Histogram Analysis Implementation Including Robust Error and Autocorrelation Estimates. *J Chem Theor Comp*, **6**(12), 3713-3720 (2010). DOI: 10.1021/ct100494z.

(20) Guixà-González, R., Rodriguez-Espigares, I., Ramírez-Anguita, J.M., Carrió-Gaspar, P., Martinez-Seara, H., Giorgino, T., & Selent, J. MEMBPLUGIN: studying membrane complexity in VMD. *Bioinformatics*, **30**(10), 1478-1480 (2014). DOI: 10.1093/bioinformatics/btu037.

(21) Yamskov, I.A., Kuskov, A.N., Babievskiĭ, K.K., Berezin, B.B., Kraiukhina, M.A., Samoĭlova, N.A., Tikhonov, V.E., & Shtil'man, M.I. New liposomal forms of antifungal antibiotics, modified by amphiphilic polymers. *Appl Biochem Microbiol*, **44**(6), 688-693 (2008).

(22) Martens-Lobenhoffer, J., Rupprecht, V., & Bode-Böger, S.M. Determination of micafungin and anidulafungin in human plasma: UV- or mass spectrometric quantification?

*J Chromatogr B Analyt Technol Biomed Life Sci*. **879**(22), 2051-2056 (2011). DOI: 10.1016/j.jchromb.2011.05.033.
